# Benchmarking of tools for resolving the plasmidome from short-read assemblies for *Klebsiella pneumoniae*

**DOI:** 10.1101/2025.07.24.666686

**Authors:** Christopher H. Connor, Ryan R Wick, Claire L. Gorrie, Mia A. Winkler, Iren H Löhr, Danielle J. Ingle, Margaret M.C. Lam

**Author notes:** Correspondence: Chris H. Connor Margaret M.C. Lam. contributed equally (Danielle J. Ingle, Margaret M.C. Lam).

## Abstract

Plasmids play a critical role in the dissemination of antimicrobial resistance genes and virulence factors in healthcare associated pathogens, such as *Klebsiella pneumoniae*. Surveillance of these plasmids relies on whole genome sequencing data often generated in clinical and public health settings which frequently use short-read platforms. Therefore, there is a need for robust, scalable tools that can identify and/or reconstruct plasmid sequences from short read data. A myriad of tools already exist to address this problem, however the optimum tool for plasmid identification in *K. pneumoniae* remains unclear. From a comprehensive search of the literature and code repositories we identified 44 plasmid identification tools, highlighting the uncertainty around best practices. Here, we sought to evaluate these 44 tools to determine which is best suited for reconstructing the plasmidome of *K. pneumoniae* and related species from the species complex (KpSC). We used a publicly available dataset of 568 diverse KpSC isolates that had both short-read Illumina data and closed hybrid assemblies available. This allowed us to investigate which tools perform best at recovering plasmid sequences when only short-read data is available, whilst knowing the ground truth. From the 44 tools, 34 were excluded as they: were intended for plasmid typing / characterisation (n=3), were not intended for KpSC (n=1), required metagenomic data (n=7), required long read data (n=1), could not be installed (n=13) or could not be run on the command line (n=10). The remaining nine tools had their precision and recall metrics calculated and combined into an overall F1 score. Each individual tool displayed the full range of F1 scores (0 to 1) across our collection of genomes, overall, the best performing was PlaScope followed closely by MOB-suite. Future tools developed in this crowded space should offer meaningful advancements over existing tools and be rigorously benchmarked using standardised datasets that reflect plasmid diversity.

**Impact Statement:** Short-read sequencing (e.g., Illumina) remains the platform of choice for routine whole-genome sequencing in many public health settings, underpinning pathogen and antimicrobial resistance (AMR) surveillance. Plasmids play a critical role in the transmission of AMR, yet accurate reconstruction of plasmids from short-read data remains a major challenge despite the availability of numerous tools. Here, we present a systematic benchmarking of plasmid reconstruction tools using a large and diverse dataset of 568 *Klebsiella pneumoniae* species complex (KpSC) genomes. Our evaluation revealed considerable variability in tool performance, with no single method capable of accurately reconstructing the entire plasmid repertoire. Based on these findings, we provide recommendations on the most suitable tools for plasmid reconstruction from short-read data for KpSC. We also outline key considerations for future tool development to improve scalability and reliability, which will be essential for strengthening genomic surveillance of plasmids and AMR.

## INTRODUCTION

*Klebsiella pneumoniae* is a leading cause of opportunistic infections in healthcare settings globally (1,2). Many of these strains (also referred to as classical *K. pneumoniae* or cKp) harbour resistance to multiple antimicrobials, resulting in treatment complications or failures (2). Widespread resistance to the third-generation cephalosporins and carbapenems has been identified as a high-priority concern by the WHO due to reliance on these last resort antimicrobials for treating infections caused by Gram-negative bacteria (3). Outside of clinical settings, *K. pneumoniae* can also cause highly invasive, community-acquired infections, often arising in otherwise healthy and immunocompetent individuals (4). These infections are typically caused by a unique subset of strains (or ‘clones’) referred to as hypervirulent *K. pneumoniae* (hvKp) that carry additional virulence determinants and are distinct from the clones that commonly cause healthcare-associated infections (2). The accessory genome varies significantly across *K. pneumoniae* clones (e.g. differentiating cKp from hvKp), and is largely comprised of mobile genetic elements, such as plasmids (2,5).

Plasmids play a key role in mediating the spread of antimicrobial resistance (AMR) genes, virulence genes, or both, in *K. pneumoniae* and closely related species (collectively referred to as the *K. pneumoniae* species complex or KpSC). Plasmids have been important drivers shaping the evolution of different clones (5). Of particular concern has been the carriage and dissemination via plasmids of AMR to third generation cephalosporins and, more recently, carbapenem classes in KpSC (6–10). There is incredible diversity in the plasmids that have been sequenced from KpSC isolates, varying in features including gene content, length and incompatibility types (5). There is also significant variation in the number and types of plasmids carried by an individual host strain and can range from 0 to upwards of 10. Interestingly, plasmid number and loads tend to be higher in cKp isolates, whereas hvKp typically carry just one large plasmid referred to as a large virulence plasmid or KpVP-1 (11). Beyond the accessory properties conferred by plasmids, plasmid transmission between strains can also potentiate clinical outbreaks (i.e. ‘plasmid outbreaks’). Given their clinical relevance, the ability to confidently identify plasmids from routinely generated whole genome sequencing (WGS) data is therefore important.

WGS data is increasingly generated and applied in routine surveillance and clinical outbreak investigations globally (12,13). Short-read sequencing Illumina platforms remain the workhorse in clinical and public health surveillance efforts (14). This is due to relatively low costs, high read accuracy, scalability and the ability to implement and accredit into validated workflows. However, a key drawback is the inability to accurately resolve large plasmids from draft genome assemblies generated from short-read sequence data, as plasmid structures generally carry repetitive sequences (15). The application of long-read sequencing on Oxford Nanopore Technologies (ONT) platforms can circumvent this issue as the reads can span across genomic repeats. As such, ONT data is commonly used in bacterial genomics to resolve complete genomes of interest, however it is not always feasible to perform long-read sequencing on all large-scale datasets as is done on Illumina platforms (16). Given the importance and interest in studying plasmids and complexities around plasmid structures, multiple tools have been developed that seek to address the challenges around accurately reconstructing or identifying plasmids from short-read-only assemblies.

At present, there are at least 44 tools that seek to identify and predict plasmid sequences, which creates the conundrum of which tool is best suited to reconstructing the plasmidome of bacterial pathogens such as those from the KpSC with complex plasmid dynamics. Tools to reconstruct plasmids from draft genome assemblies continue to be developed, with at least eight tools developed since the initiation of this study in June 2023. The majority of these tools will either predict whether a contig from an input genome assembly is plasmid or chromosomal sequence (i.e. binary classifier) or attempt to reconstruct the plasmids from input reads (i.e. *de novo* plasmid assemblers). Given the growing number of available tools, it is difficult to know which to use, particularly for KpSC isolates where the plasmid landscape is diverse and complex.

There have been very few independent evaluations performed solely to assess tool performance, beyond initial benchmarking undertaken in publications linked to different tools (15,17,18). The first of these was a species-agnostic study comparing four automated plasmid prediction tools (plasmidSPAdes (19), Recycler (20), cBar (21) and PlasmidFinder (22)) across 42 genomes from 12 bacterial genera that included 12 *K. pneumoniae* genomes (15). The second was *E. coli* focussed and benchmarked six tools (plasmidSPAdes (19), HyAsP (23), FishingForPlasmids (15), SCAPP (24), MOB-suite (25) and gplas (26)) using a diverse collection of 240 *E. coli* genomes (17). The third used a dataset of 93 Enterobacterales genomes (from 14 species) and 157 *Enterococcus* species and benchmarked 13 tools including Plasmer, PlasmidEC, PlaScope, and gplas2 (18,27–29). All these evaluations concordantly reported that none of the tools accurately recovered their study’s entire plasmid dataset but differed in which tools performed best for their respective datasets. Arredondo-Alonso *et al* identified that all tools’ performance was generally improved for smaller plasmids defined as <50 Kbp, and that while PlasmidFinder had the best predictions it frequently missed plasmids (15). In contrast, Paganini *et al* identified that the majority of plasmids could not be correctly resolved by any of the tools, with MOB-suite performing best for plasmid recovery and plasmidSPAdes best for recovering small plasmids <18 Kbp in size (17). Finally, Teixeira *et al* found that PlaScope performed well for the Enterobacterales at plasmid detection, while gplas2 was best for plasmid reconstruction, with a limitation being that the 93 Enterobacterales genomes were more likely to have AMR genes carried on plasmids that resulted in shorter contigs that were difficult to classify (18). Ultimately the optimum tool for plasmid identification in KpSC remains unclear.

Here, we sought to evaluate the tools that met our inclusion criteria to determine which was best suited for reconstructing the plasmidome of KpSC genomes. We leveraged a well-characterised dataset of 568 KpSC isolates with high-quality Illumina and matched ONT hybrid assemblies, and plasmid content that reflects global KpSC plasmid diversity (31). This allowed us to address the question of when only short-read data is available for large-scale datasets of bacterial species with diverse plasmid content, which tool is best to identify and predict plasmid sequences.

## METHODS

### Tool identification and selection criteria for inclusion in benchmarking

We first performed a search of the literature for available plasmid prediction tools (5^th^ January 2026) by using the same NCBI PubMed search parameters that were specified in the *E. coli* study (17) (**Supplementary Material**). The resulting publications were manually processed to identify tools for inclusion in this study. To capture tools that had not been linked to a publication, we additionally performed an extensive internet search alongside searches of commonly used code repositories (GitHub, Bitbucket, GitLab) using the search terms “plasmid” and “plasmid detection”. Lastly, we reviewed previous benchmarking publications to ensure no other tools had been missed. This generated a comprehensive list of relevant plasmid tools (**Supplementary Table 1**).

We then proceeded to identify the relevant tools to benchmark for their abilities to reconstruct plasmids from short-read data. Several selection criteria were applied to be included in this study. Specifically, the tool must:

1. predict, identify or assemble plasmid sequences (i.e. generate plasmid sequences, not just plasmid typing or gene information),
2. accept as input short-read-only data as either reads, genome assemblies or assembly graphs,
3. detect plasmids in data sequenced from a single genome (i.e. not a metagenomic sample),
4. run on the command line,
5. be successfully installed via the command line with clear instructions on installation & usage, and not require proprietary software,
6. run without encountering an error with default parameters.

In the event where a tool had failed to install, a second attempt at installation was carried out on an alternative computer. A minimum level of troubleshooting was performed. For example where a specific dependency was reported as missing, the dependency was installed and installation reattempted. Further, if a tool did not immediately run with default parameters, a second installation and test run on an alternative device was performed. Details, such as installation commands, error messages and troubleshooting steps for the tools which could not be installed or run, were documented and are available at github.com/C-Connor/PlasmidToolBenchmarking/blob/main/ToolInstallations.md.

### Benchmarking dataset

The dataset used for benchmarking the tools that passed the selection criteria consisted of 568 KpSC genomes from a recently published study (30). These genomes belonged to diverse lineages within *K. pneumoniae* (n=482) and the broader species complex, which included 69 *K. variicola*, nine *K. quasipneumoniae* subspecies *similipneumoniae*, seven *K. quasipneumoniae* subspecies *quasipneumoniae* and one *K. quasivariicola.* These genomes are a subset from a larger study collection of >3000 Illumina-whole genome sequenced isolates collected in Norway between 2001 to 2020 from human, animal and marine samples (32). Importantly, in addition to Illumina short-read sequencing, all 568 isolates also had ONT long-read data available which had been assembled into complete genomes. These complete genomes provided the ground truth for assessing the performance of each tool. The accessions, assembly stats and sequencing details (i.e. Illumina platform, ONT flow cell kits) can be found in Table 1 linked with the original publication by Hetland *et al* (figshare: <u>10.6084/m9.figshare.c.7622345</u>) (32).

**Table 1.**
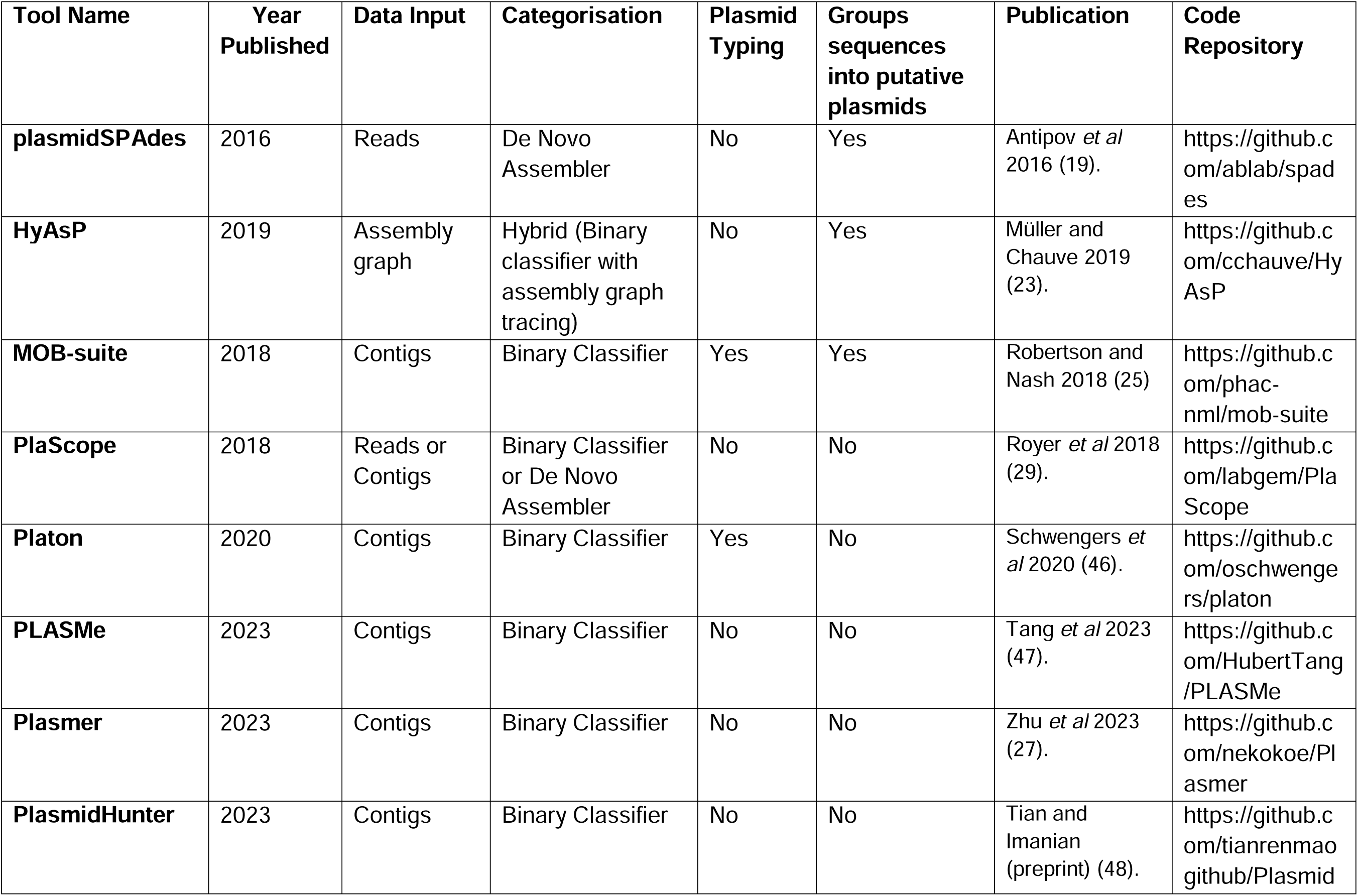
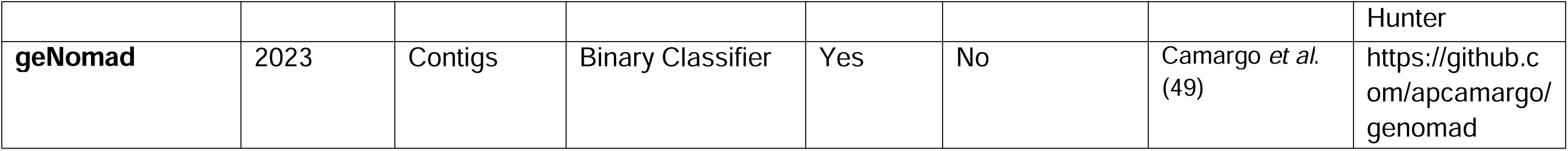
Tools benchmarked in this study.

| **Tool Name** | **Year Published** | **Data Input** | **Categorisation** | **Plasmid Typing** | **Groups sequences into putative plasmids** | **Publication** | **Code Repository** |
| --- | --- | --- | --- | --- | --- | --- | --- |
| **plasmidSPAdes** | 2016 | Reads | De Novo Assembler | No | Yes | Antipov *et al* 2016 (19). | https://github.com/ablab/spades |
| **HyAsP** | 2019 | Assembly graph | Hybrid (Binary classifier with assembly graph tracing) | No | Yes | Müller and Chauve 2019 (23). | https://github.com/cchauve/HyAsP |
| **MOB-suite** | 2018 | Contigs | Binary Classifier | Yes | Yes | Robertson and Nash 2018 (25) | https://github.com/phac-nml/mob-suite |
| **PlaScope** | 2018 | Reads or Contigs | Binary Classifier or De Novo Assembler | No | No | Royer *et al* 2018 (29). | https://github.com/labgem/PlaScope |
| **Platon** | 2020 | Contigs | Binary Classifier | Yes | No | Schwengers *et al* 2020 (46). | https://github.com/oschwengers/platon |
| **PLASMe** | 2023 | Contigs | Binary Classifier | No | No | Tang *et al* 2023 (47). | https://github.com/HubertTang/PLASMe |
| **Plasmer** | 2023 | Contigs | Binary Classifier | No | No | Zhu *et al* 2023 (27). | https://github.com/nekokoe/Plasmer |
| **PlasmidHunter** | 2023 | Contigs | Binary Classifier | No | No | Tian and Imanian (preprint) (48). | https://github.com/tianrenmao/github/Plasmid |
|  |  |  |  |  |  |  | Hunter |
| **geNomad** | 2023 | Contigs | Binary Classifier | Yes | No | Camargo *et al.* (49) | https://github.com/apcamargo/genomad |

### Benchmarking of selected plasmid tools

Benchmarking for the tools that passed our selection criteria was implemented as part of a Nextflow pipeline (github.com/C-Connor/PlasmidToolBenchmarking), and an overview of this pipeline is provided in **Figure 1** and **Supplementary Figure 1**. Compute time and RAM usage for each tool was recorded and collated through the Nextflow trace file. Each tool was run using its default parameters and was allocated 8 CPUs through Nextflow, including tools that do not provide multi-threading support (MOB-suite). The input data for each tool is detailed in **Table 1**. Genome assemblies, or assembly graphs, were generated using SPAdes v3.15.4 and Unicycler v0.5.1 (33,34). We chose to assemble our short-read data with SPAdes as it is widely used in the field, particularly in public health settings. Unicycler was chosen as a comparator assembler as while it uses SPAdes it has several improvements, including automatic *k*-mer size selection, overlap removal and a contamination filter (34). To assess the accuracy of the predictions for each tool, the outputs were compared to the completed hybrid genome assemblies (30) using BLASTn v2.16.0 (35), whereby the query and subject corresponded to the tool outputs and the complete genomes of each sample, respectively (**Supplementary Figure 1**). A high stringency filter was applied of 80% sequence identity and 80% query coverage; these thresholds are commonly used in the field (36).

**Figure 1.**
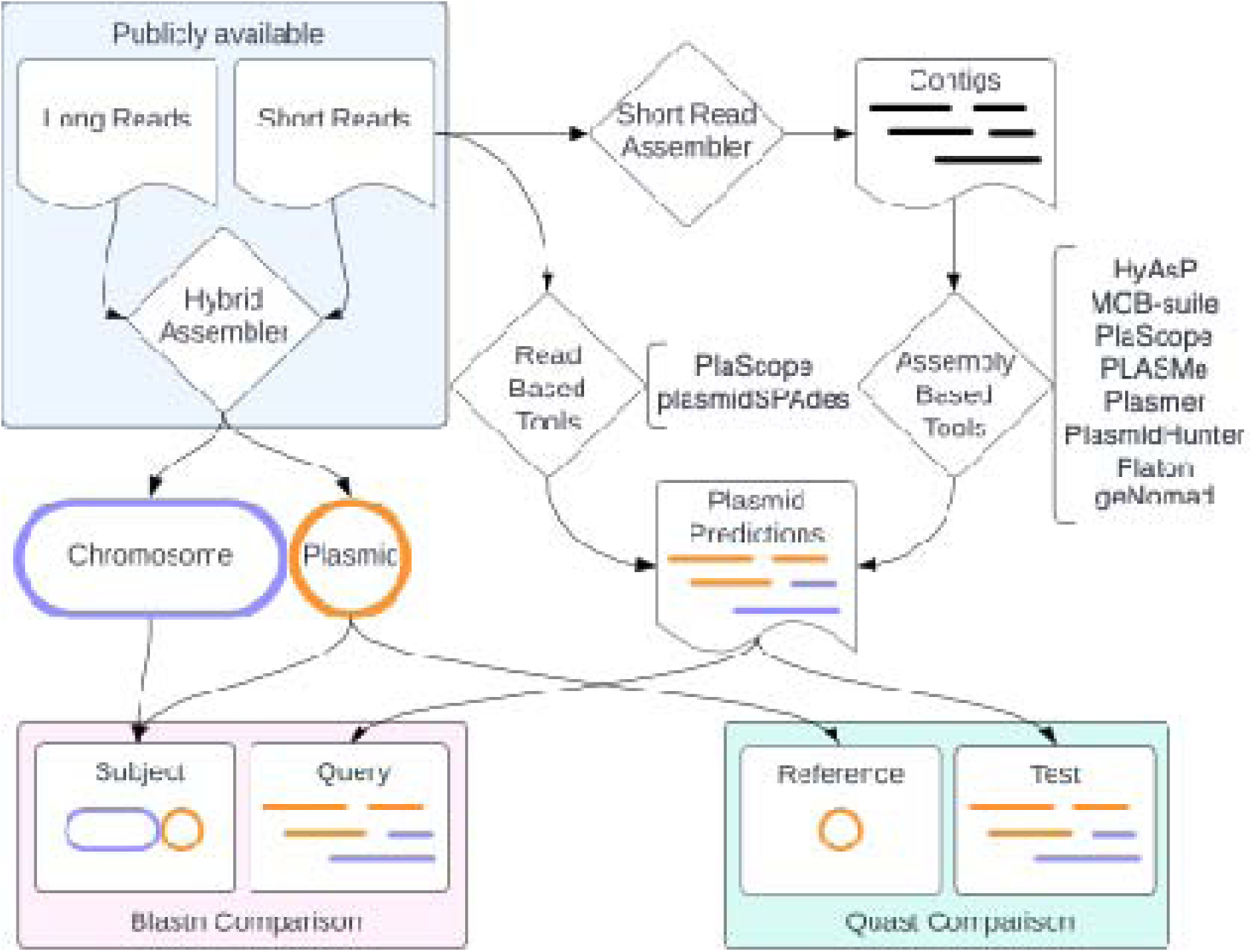
Schematic depicting the data flow for analysis. Publicly available data taken from Hetland *et al.* (30) in the form of complete hybrid assemblies and short-read-only data. Short reads were assembled into short-read-only assemblies and used as input for plasmid prediction tools. Predictions from the plasmid tools were assessed against the complete hybrid assemblies using BLASTn and Quast. The BLASTn approach was first applied to initially quantify rates of contig identification and generate true positive / negative and false positive / negative rates. Quast was subsequently used to quantify precision, recall and F1 scores.

To collect more detailed metrics around output accuracy, we used the genome assembly evaluation tool Quast v5.3.0 (37). The plasmid predictions were compared against the total plasmid content of each sample in addition to the individual plasmid sequences from the complete genomes (**Figure 1** and **Supplementary Figure 1**). Post-run processing of the tool outputs (i.e. to convert outputs into a uniform format for analysis) and assessment of accuracy via BLASTn and Quast comparisons were both implemented as part of the Nextflow pipeline (available at: github.com/C-Connor/PlasmidToolBenchmarking). Using the metrics from Quast, we calculated three key metrics for assessing performance, and a schematic of how these metrics were calculated and what they represent is provided in **Supplementary Figure 1**.

These were:

1. Total plasmid content precision, calculated by dividing the total length of plasmid predictions that could be aligned to a reference plasmid by the total length of all plasmid predictions. This metric represents what proportion of the plasmid sequences predicted matched to a plasmid in the hybrid genomes.
2. Total plasmid content recall (the genome fraction metric from Quast), represents the proportion of the reference plasmid sequence (from the hybrid genomes) that is aligned to a contig in the plasmid prediction collection,
3. Individual plasmid recall (genome fraction metric from Quast) for individual plasmids in the hybrid genomes
4. A summary F1 score for total plasmid content calculated as (2 x Precision x Recall / (Precision + Recall)). (17)

The precision metric, and hence the summary F1 score, could not be calculated at the individual plasmid level as most tools do not divide their plasmid predictions into individual putative plasmids. This makes calculating the denominator for precision (total plasmid predictions) challenging as it is unclear which plasmid bin incorrect (chromosome) sequences should be assigned (**Supplementary Figure 1**). As F1 scores are bounded between 0 and 1 and not normally distributed, statistically significant differences in F1 score were tested with pairwise comparisons using the non-parametric Wilcoxon signed rank test with continuity correction and Bonferroni p value adjustment. Pairs of tools that were found to be statistically different had their per sample F1 scores compared to calculate the median difference in F1 score (effect size). Statistical tests were performed in R v4.5.0.

### Dataset metrics and typing

Additional metrics and isolate typing were performed on the benchmarking dataset to correlate with plasmid tool performance. Short-read-only assembly quality was measured using Quast supplying the hybrid assembly for each isolate as the ‘reference’ genome. A count of total contigs and their sizes in each short-read-only assembly was produced by SeqKit (38). AMR genes were detected with Kleborate v3.2.4 using the ‘kpsc’ modules (39). The count of IS elements in each plasmid were obtained from Winkler *et al* supplementary materials (31). Dead ends in the assembly graph were quantified with GFA-dead-end-counter (40). Sequencing depth was quantified by first mapping the short reads to the hybrid assembly with minimap2 v2.30 using the short-read preset (41). Depth was then calculated from the read alignment using Samtools coverage v1.23 (42). Sharing of *k*-mers between the chromosome and plasmids was quantified with Mash v2.3 using the hybrid genomes (43). Chromosome and plasmid contigs were separated with SeqKit and individually sketched with a sketch size of 100,000. Sharing of *k*-mers was then calculated with Mash distance. Results from typing metrics were collated with CSVTK v0.31.0 (44) and Python v3.13.3. Associations between tool performance (precision, recall, F1) and additional metrics were tested using Spearman’s Rank Correlation test in R. All code for calculating metrics and typing is included in the Nextflow pipeline and associated R code, and both are available at github.com/C-Connor/PlasmidToolBenchmarking. Result files for all the data presented in this manuscript are also available on the GitHub.

## Data visualization

Performance metrics and graphs were made in R v4.5.0 with ggplot2 v3.5.2 (45). The code for reproducing the figures is available at github.com/C-Connor/PlasmidToolBenchmarking.

## RESULTS AND DISCUSSION

### Multiple tools developed to tackle the problem of plasmids

Plasmid genomics is a rapidly growing area of microbial research, underpinned by advances in sequencing technologies and interest in resolving plasmids and their gene content as they drive population expansions that can lead to outbreaks and the spread of AMR (**Figure 2A**). We identified a total of 44 plasmid detection or reconstruction tools, 42 of which have been released since 2015 (**Figure 2B**). This demonstrates that there is active interest in trying to develop tools suited to identifying plasmids in bacterial genomics datasets. However, after the application of our selection criteria (see Methods), only eight tools were included in the full benchmarking analyses (**Figure 2C**).

**Figure 2.**
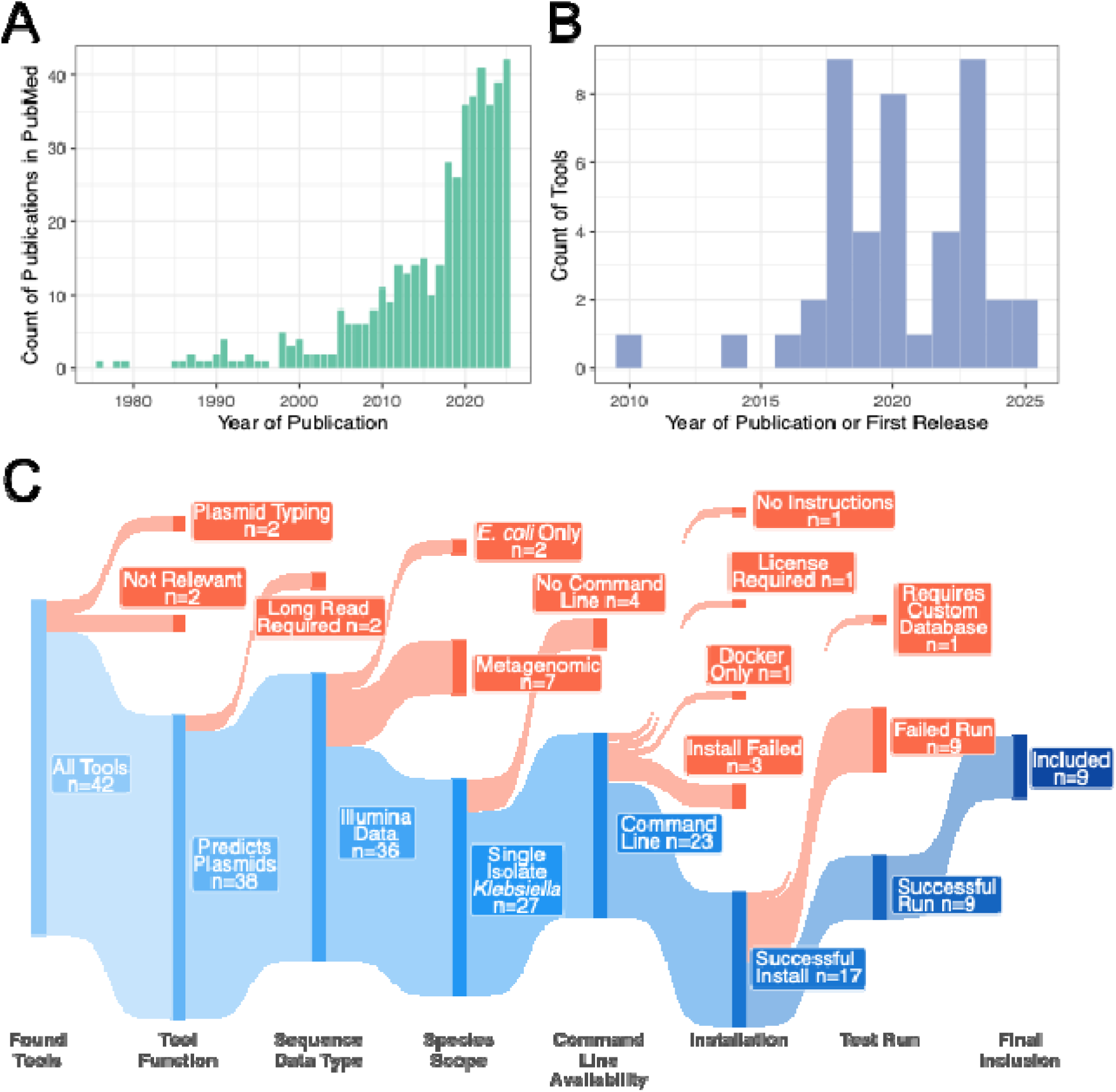
Identification and selection of plasmid tools for benchmarking analysis. Panel A) Bar plot of publications matching our PubMed search criteria over time (years). Panel B) Bar plot of 44 tools (details in **Supplementary Table** 1) plotted over time with year of publication, where the tool has an associated publication, otherwise year of first release is used. Panel C) Overview of inclusion criteria and exclusion reasons for the 44 plasmid identification tools.

Tools were excluded at all stages of our selection criteria (**Figure 2C**). Two tools were excluded as they did not produce full plasmid sequence predictions (PlasmidFinder and PlasmidSeeker), instead providing the user with plasmid associated genes. These two tools were classed as plasmid typing tools. One tool was excluded as it did not support identification of plasmid sequences and instead focussed on identifying DNA sequences associated with a specific phenotypic trait and was classed as ‘not relevant’ (PlasmidTron). The gplas2 tool was excluded as it required plasmid predictions to be performed by a separate tool before attempting to isolate individual plasmids, the authors recommend the use of plasmidEC as input however any binary classifier is accepted. Of the remaining 40 tools, 38 were short-read data compatible, taking as input either raw FASTQ files, genome assemblies in FASTA format or assembly graphs. We did not consider tools where long-read data was required (plasmIDent and plassembler) as we sought to explore only tools that would suit routinely generated short-read data. A further seven tools were subsequently excluded at the *Species Scope* stage as they were intended for use with metagenomic datasets (metaplasmidSPAdes, Plasmid_Assembler / plasmidextractor, PlasmidPicker, cBar, PPR-Meta, PlasGUN, and 3CAC), which was not an objective of this study, and another two tools were excluded because they were specific for *E. coli* datasets (PlasmidEC and FishingForPlasmids) (**Figure 2C**).

Tools that did not explicitly state which organisms they could support were assumed to be applicable to KpSC genomes. Of the 29 remaining tools that were suitable for detecting plasmids from KpSC genome datasets, four were excluded at the *Command Line Availability* stage as there were no available command line versions of their tools (**Figure 2C**). The rationale for this decision was that command line use is optimal for working with larger datasets. The *Installation* stage excluded six tools. Of those six tools, one (Deeplasmid) was excluded as it was only available as a Docker container, one (PlasBin-flow) as it required a proprietary licence, and one (GraphPlas) because it provided no installation or usage instructions. The remaining three tools all encountered errors during their installation (see github.com/C-Connor/PlasmidToolBenchmarking/blob/main/ToolInstallations.md for details on install commands and error messages). A total of 19 tools were installed however nine tools encountered an error when performing a test run and so were excluded. An additional tool was excluded (ACCIO) as it required a custom database that was not provided. The remaining nine tools were used for the benchmarking analysis: geNomad, HyAsP, plasmidSPAdes, MOB-suite, PlaScope, Platon, PLASMe, Plasmer, and PlasmidHunter (19,23,25,27,29,46–49) (**Figure 2C**, **Table 1**). Of special note only two of the included tools (MOB-suite and HyAsP) create individual plasmid sequence files. All other included tools instead produce a single plasmid output file representing the total plasmid content of the isolate (plasmidome).

The high number of plasmid detection tools obscures the fact that many are unsuitable for modern large-scale bacterial datasets. The reduction from 44 identified tools to nine for our benchmarking purposes is substantial but unsurprising, particularly given the prevalence of ‘abandonware’ in bioinformatics. Only eight of the 44 tools had received code updates since 2025, while 14 had not been updated in over five years. This pattern of tool abandonment likely reflects the lack of academic incentives for ongoing software maintenance (50). Of the nine tools benchmarked in this study, two (geNomad and Platon) had been updated within the past 12 months, three had been updated within the past 2 years (MOB-suite, PLASMe and PlasmidHunter), with the remaining 4 tools (plasmidSPAdes, Plasmer, HyAsP and PlaScope) receiving updates within the past 3 to 5 years.

### Assessing runtime and RAM usage

The plasmid tools that met our inclusion criteria were run on our test dataset using a Nextflow pipeline (available at: github.com/C-Connor/PlasmidToolBenchmarking), which recorded analysis time and compute resource usage. The nine tools were split into two broad groups reflecting functionality of the tool, namely assembly-based tools and read-based tools. Assembly-based tools (also referred to as binary classifiers) predict if each contig from a draft genome assembly is plasmid or chromosome. While read-based tools (also referred to as *de novo* assemblers) accept sequencing reads to reconstruct plasmid sequences. Of special note are the benchmarked tools PlaScope and HyAsP, whereby the former acts as a binary classifier but can accept both reads and assemblies so was assessed in both modes, while HyAsP requires assembly graphs and uses a hybrid plasmid identification approach (combining binary classification with *de novo* assembly methods to trace paths through the graph) (23,29).

PlaScope (assembly input) demonstrated the fastest performance with a median runtime of 4.8 seconds but also produced the slowest median runtime of all tools (23.96 minutes) when given reads as input (**Figure 3A**). In general, read-based tools were slower than assembly-based tools and showed the greatest variability in runtime (PlaScope (reads) IQR: 18.13 to 37.24 minutes, plasmidSPAdes IQR: 13.93 to 30.18 minutes) (**Figure 3A**). However, the runtime for the assembly-based tools does not consider the time required to generate the genome assemblies, which with SPAdes was on average (median) 17.89 minutes and with Unicycler was 51.53 minutes (**Figure 3A**). Factoring this additional assembly time for the assembly-based tools would in fact make many of the runtimes across various tools comparable.

**Figure 3.**
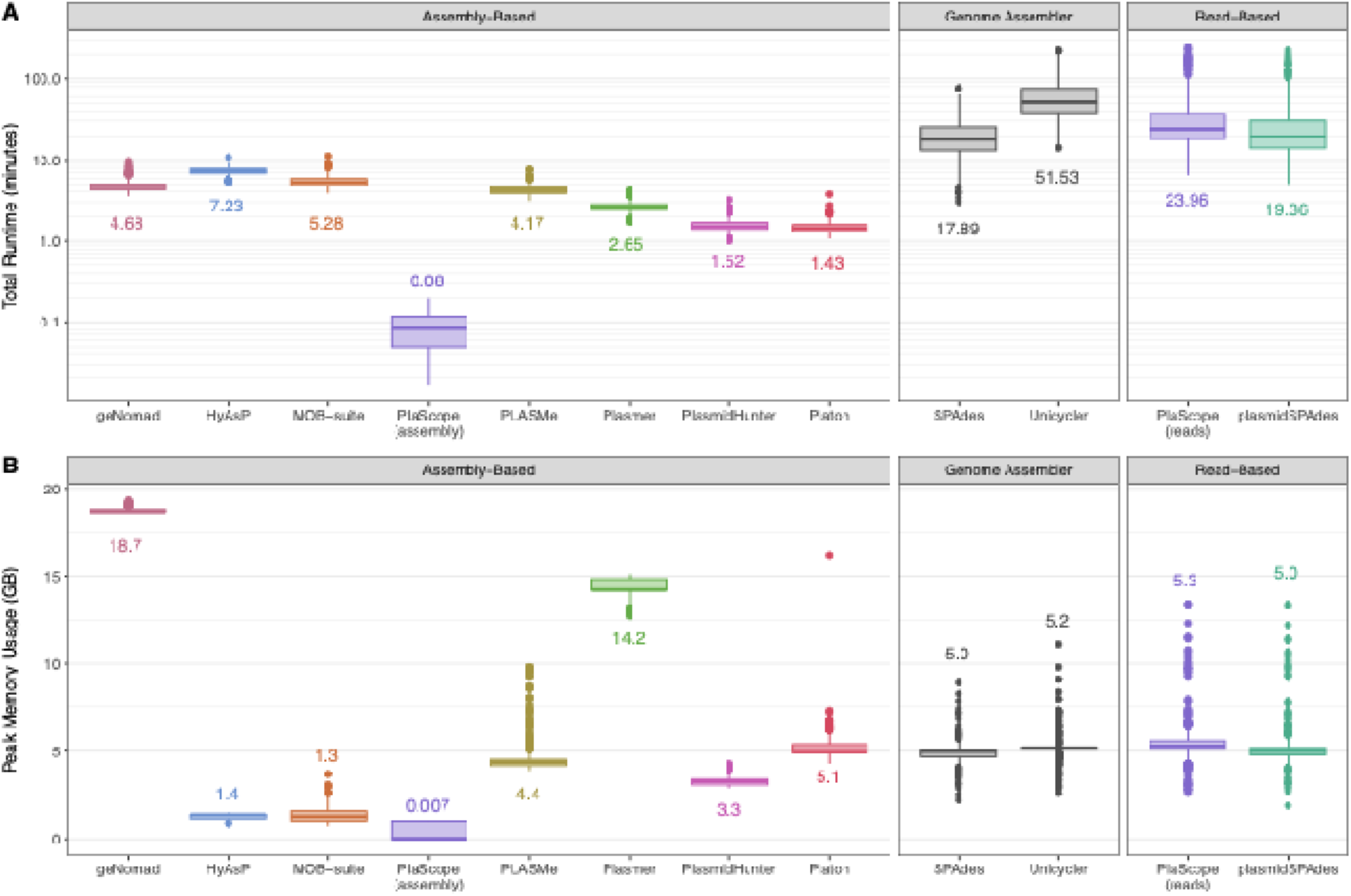
Plasmid tool runtimes and memory usage Panel A) Wall-clock time taken for plasmid prediction tools to run in minutes (log scale). Median runtimes are labelled. Panel B) Peak memory usage (RAM) in gigabytes for each tool. Median peak memory usage for each tool is labelled.

**Figure 4.**
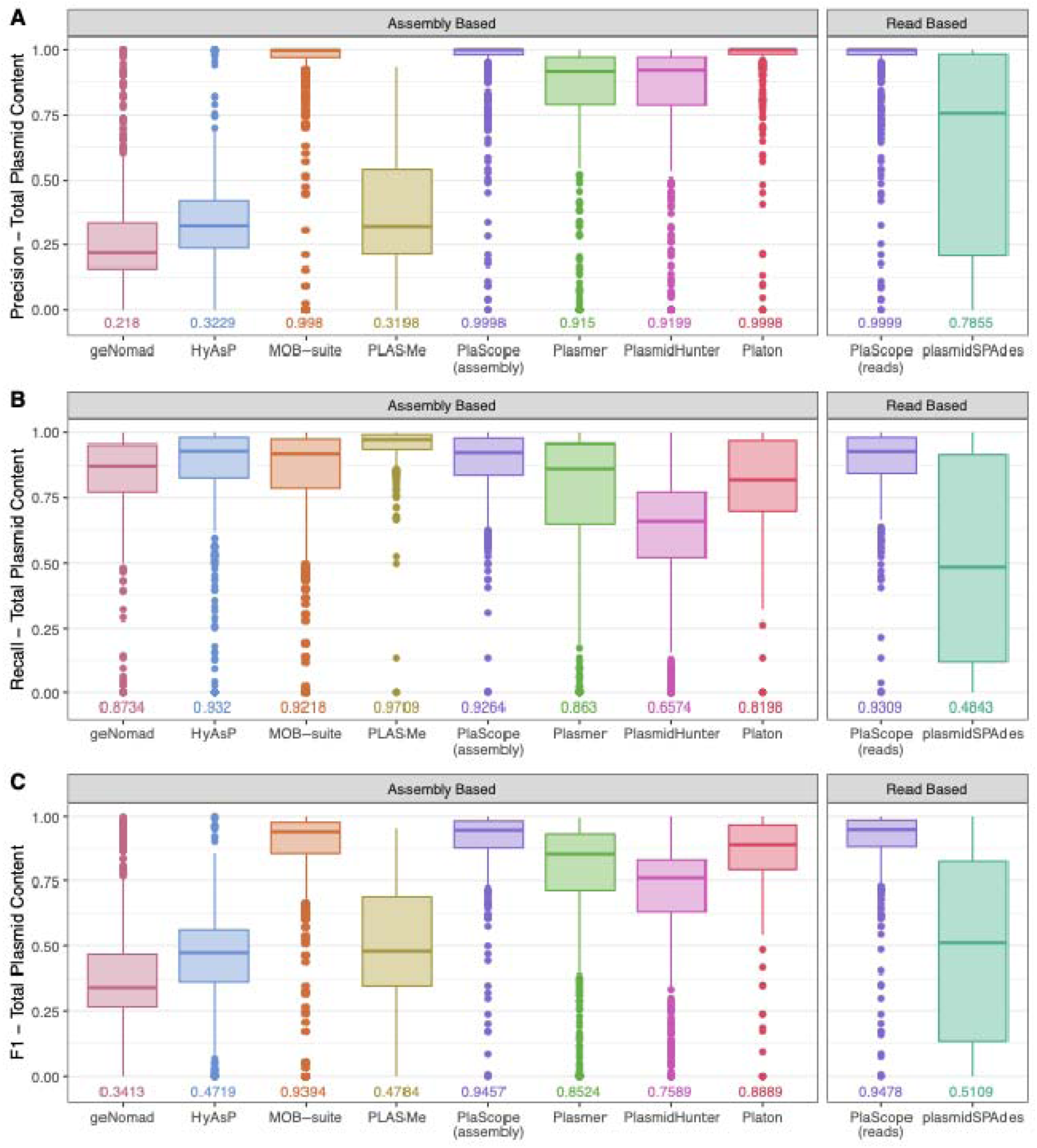
Assessing precision, recall and F1 scores for the benchmarked tools. Panel A) Boxplot of the precision score for each tool. The tool precision represents the proportion of plasmid predictions that matched plasmid sequences, whereby a value of 1.0 indicates all predictions were true plasmids while 0.0 indicates all predictions were chromosomal sequences. Median precision scores for the tools are labelled. Panel B) Boxplot of the recall score for each tool. The recall represents whether the tool was able to identify all the plasmid sequences present in the sample. A recall of 1.0 means the tool was able to identify all the plasmid sequences present, while 0.0 means all plasmid sequences in the sample were missed. Median recall scores for the tools are labelled. Panel C) Boxplot of the overall F1 score for each tool, calculated from precision and recall scores. The median F1 scores for the tools are labelled.

Except for geNomad and Plasmer, peak memory usage was generally low, with most tools using less than 5 GB (**Figure 3B**). geNomad had the highest average peak usage at 18.7 GB followed by Plasmer at 14.2 GB, while PlaScope (assembly input) had the lowest memory requirement with an average of 14 MB peak usage. Depending on compute resources, the higher RAM usage by geNomad may be a cause to select alternative assembly-based tools. The two read-based tools, PlaScope and plasmidSPAdes had comparable RAM usage, similar to that of the genome assemblers (SPAdes and Unicycler).

### Overview of benchmarking dataset

The KpSC benchmarking dataset of complete hybrid genomes comprised 568 chromosomes and 1,381 complete plasmids (30). Plasmids were present in 504/568 genomes and demonstrated variability in number, size and association with AMR and/or virulence, providing a globally representative large-scale dataset to investigate the plasmidome (31). Plasmids present in this dataset have been confirmed to represent global plasmid diversity not just from clinical but also environmental sources (31). The remaining genomes with no plasmids (n=64) were included in our analysis as a negative control. The median number of plasmids per sample was 2, with a range of 1 to 10 across the 504 genomes that carried plasmids. Plasmid lengths ranged from 1,240 bp to 424,266 bp, highlighting significant diversity in plasmid carriage and types within the entire collection.

Across the dataset of short-read-only SPAdes assemblies there were a total of 264,105 contigs, of which 55,228 were longer than 500 bp in length (**Supplementary Figure 2**). The contigs from these fragmented assemblies were classified as chromosomal or plasmid-derived using BLASTn against the complete genomes as the reference sequences. Only high-confidence matches (i.e. those defined as >80% identity, >80% query coverage and consistent match to chromosome or plasmid) were retained for further analysis (**Supplementary Figure 1 & 3)**.

Based on these thresholds, 117,619 contigs (44.5%) were confidently assigned, comprising 93,896 chromosomal and 23,723 plasmid-derived contigs, with a collective length of 3,002 Mbp and 112.1 Mbp, respectively.

### Assessing performance to reconstruct the plasmidome of *Klebsiella*

We next explored the ability of the nine tools to predict plasmids in short-read data. Consistent with previous studies, none of the tools were able to correctly identify all 23,723 contigs that were derived from plasmid sequences (**Table 2, Supplementary Figure 3**). PLASMe had the highest number of plasmid predictions (67,445) and identified 22,247 of 23,723 plasmid contigs (93.8% recovery rate), however PLASMe also reported 45,198 contigs as putative plasmid matches but BLASTn matched them to the chromosome. This corresponds to a true positive rate of only 33.0%, which was the lowest across all benchmarked tools (**Table 2, Supplementary Figure 3**). Platon had the highest true positive rate with 92.9% of the 5,611 plasmid predictions matching to plasmid sequences in the hybrid genomes. Platon was closely followed by PlaScope (assembly input) with 92.7%. However, Platon and PlaScope (assembly input) were only able to recover a fraction of the total plasmid content of the isolates, recovering 23.7% and 31.1% respectively of all BLASTn plasmid matches. The recovery rate of Platon was the lowest of all benchmarked tools. These results highlight trade-offs between prediction accuracy and sensitivity when trying to reconstruct the plasmidome of *Klebsiella* and pose a dilemma for users when considering tool selection. For example, Platon yields higher confidence in plasmid predictions but risks missing important plasmids, while PLASMe facilitates more comprehensive detection but at the cost of a high rate of false positives (i.e. chromosomal sequences being flagged as plasmid). Benchmarking studies, such as this large *Klebsiella* dataset, provide important data that can inform the selection of tools best suited to public health surveillance or research settings.

**Table 2.** Contig classifications for benchmarked tools grouped by BLASTn match to hybrid assemblies.

| Tool | Data Input | Contig Predicted as Plasmid by Tool |  | Contig Predicted as Chromosome by Tool |  | Contig Not Classified by Tool |  | Total |
| --- | --- | --- | --- | --- | --- | --- | --- | --- |
|  |  | BLASTn Match to Plasmid | BLASTn Match to Chromosome | BLASTn Match to Plasmid | BLASTn Match to Chromosome | BLASTn Match to Plasmid | BLASTn Match to Chromosome |  |
|  |  | True Positive | False Positive | False Negative | True Negative |  |  |  |
| **geNomad** | **SPAdes Assembly** | 5867 | 5165 | 0 | 0 | 17856 | 88731 | 117619 |
| **HyAsP** | **SPAdes Assembly graph** | 5328 | 2814 | 0 | 0 | 0 | 0 | 8142 |
| **MOB-suite** | **SPAdes Assembly** | 6195 | 1115 | 17528 | 92781 | 0 | 0 | 117619 |
| **PlaScope (assembly)** | **SPAdes Assembly** | 7383 | 578 | 300 | 29574 | 16040 | 63744 | 117619 |
| **PLASMe** | **SPAdes Assembly** | 22247 | 45198 | 0 | 0 | 1476 | 48698 | 117619 |
| **Plasmer** | **SPAdes Assembly** | 7463 | 4138 | 1426 | 30513 | 14834 | 59245 | 117619 |
| **PlasmidHunter** | **SPAdes Assembly** | 7225 | 3236 | 151 | 23401 | 16347 | 67259 | 117619 |
| **Platon** | **SPAdes Assembly** | 5611 | 426 | 18112 | 85756 | 0 | 7714 | 117619 |
| **PlaScope (reads)** | **Reads** | 6967 | 597 | 286 | 27256 | 0 | 0 | 35106 |
| **plasmidSPAdes** | **Reads** | 7106 | 5947 | 0 | 0 | 0 | 0 | 13053 |

To further investigate tool accuracy, we used Quast to compare the plasmid predictions against the complete genome references. Unlike the binary insights provided from BLASTn, this approach allowed for more detailed investigation of the resolution of individual plasmids and complex sequence rearrangements (**Figure 1, Supplementary Figure 1**). Quast metrics were used to calculate precision, recall and an F1 score. Precision represents the proportion (calculated in base pairs) of plasmid sequence predictions that could be mapped to a plasmid in the complete genome reference. A precision value of 1.0 means that all predicted sequences match to plasmid sequences, while 0.0 means none of the predictions matched to plasmids (chromosome contamination). The highest median precision scores were achieved by PlaScope (read input mode) with a score of 0.9999, closely followed by PlaScope (assembly input mode) and Platon, both with a score of 0.9998. geNomad recorded the lowest median precision score at 0.218.

The recall metric represents what fraction of plasmid sequences that have been recovered by the tool, whereby a value of 1.0 means the full length of plasmid sequences have been identified, while 0.0 means the plasmid sequence is completely absent. The highest median recall score was achieved by PLASMe with 0.9709 followed by HyAsP at 0.932. PlaScope scored 0.9309 and 0.9264 for read input and assembly input respectively. While our BLASTn analysis indicated that PlaScope was only detecting 26.5% of plasmid contigs, Quast revealed that most plasmid sequences had been recovered (recall scores of 0.9309 and 0.9264), suggesting the absent contigs were small. The precision and recall scores calculated were combined into an overall F1 score, with values closer to 1.0 indicating better performance (**Figure 1, Supplementary Figure 1**). The best performing tool was PlaScope scoring a median F1 score of 0.9478 in read input mode and 0.9457 in assembly input mode. PlaScope was followed by MOB-suite with a median F1 score of 0.9394 and Platon at 0.8889. While PlaScope performed significantly better than MOB-suite (Wilcoxon signed rank p<0.005), the median difference in F1 score was only 0.002 (**Supplementary Table 2**). The tools with the weakest overall performance (lowest median F1 score) were geNomad at 0.3413 and HyAsP at 0.4719.

### Choice of genome assembler

We performed initial benchmarking using SPAdes as our genome assembler as it is widely used in the microbial genomics field. We investigated whether changing genome assembler to Unicycler would impact our results for our assembly-based tools. While Unicycler is based on SPAdes, it implements additional and more conservative assembly steps that can result in improved genome assemblies (34). Assemblies produced by Unicycler were of higher quality with fewer short contigs and a higher auNGA (a metric of assembly quality, see lh3.github.io/2020/04/08/a-new-metric-on-assembly-contiguity, **Supplementary Figure 2**).

SPAdes had produced 266,054 contigs across the 568 isolates while Unicycler only produced 73,704, of which 39,655 were over 500bp in length. Repeating our BLASTn analysis assigned 71,224 of the Unicycler contigs to hybrid assembly chromosome sequences (n=56,166) or plasmid sequences (n=15,058), giving an identification rate of 95.3% that was higher than SPAdes (44.5%). Our BLASTn analysis found that PLASMe still had the highest recovery of plasmid contigs, increasing from 93.8% with SPAdes to 99.0% for Unicycler (**Supplementary Table 3**), and also retained the worst true positive rate at 36.0% of predictions being correct. Using our Quast method to quantify recall, precision and F1 scores had shown that PlaScope achieved the best score, however when provided with Unicycler assemblies it failed to make any predictions with all contigs returned as unclassified. There are several issues on the code repository that highlight this problem (github.com/labgem/PlaScope/issues). The code base has been updated to allow support for Unicycler assemblies however this code has not been packaged into a release and so users following the installation instructions (conda based) will install an older version. Users will only be able to analyse SPAdes assemblies, and they cannot modify the FASTA headers of their SPAdes input assembly. This, again, highlights the ongoing requirement for bioinformatic software to be actively maintained throughout its lifetime, with minimal resourcing and benefit to developers. The best performing tool with Unicycler assemblies was MOB-suite with an F1 score of 0.9218, slightly reduced from the 0.9394 score with SPAdes assembly input (**Supplementary Figure 4**).

While little difference in plasmid tool performance was observed when switching from SPAdes to Unicycler, the quality of genome assemblies did increase (**Supplementary Figure 2)**. This was most noticeable from the duplication metric produced by Quast. This metric represents the level of duplication present in the genome assembly, with values over 1 indicating that sequences have been repeated and values less than 1 indicating small deletion events.

Duplication occurring in the genome assembly step can feed forward into the plasmid prediction tools and was most evident in short plasmids (**Supplementary Figure 5**). In some severe cases the duplication ratio was over 2 with SPAdes, indicating that the plasmid sequence had more than doubled. Switching the input assembler to Unicycler reduced the majority of duplication ratios to below 1.5. We confirmed that this was a result of the genome assembly input as the duplication ratio in the input assembly mirrored the duplication of plasmid predictions (**Supplementary Figure 6**).

### Impact of plasmid length, AMR and IS elements on performance

After investigating genome assembler impact on performance, we examined whether plasmid characteristics could impact performance such as plasmid length, carriage of AMR genes, prevalence of insertion sequence (IS) elements or *k*-mer sharing between plasmids and the chromosome. Associations between tools performance and plasmid characteristics were tested using Spearman’s rank correlation test (r_s_), a non-parametric test suited for bounded data (e.g. precision, recall and F1 scores) (**Supplementary Table 4**). Values for Spearman’s test range from −1 to 1, where −1 indicates a complete negative correlation, 0 indicates no correlation and 1 a complete positive correlation. The test can identify linear and non-linear associations.

We first examined how total plasmid length (total length of all plasmids present in a single isolate, calculated from the hybrid assemblies) correlated with F1 score, which revealed minimal correlation (MOB-suite r_s_=-0.20, PlaScope (assembly) r_s_=-0.15, PlaScope (reads) r_s_=-0.18) (**Supplementary Figure 7**). For the highest performing tools (PlaScope and MOB-suite) a total plasmid length of less than 100kbp generally results in an F1 score of 1.0. Isolates with a total plasmid length above 100kbp showed a high degree of variability in performance ranging from 0.75 to 1.0, with no clear correlation. Investigating the length of individual plasmid sequences against recall score (a combined F1 score cannot be calculated at the individual plasmid level, **Supplementary Figure 1**) confirmed that longer plasmids had worse performance (MOB-suite r_s_=-0.37, PlaScope (assembly) r_s_=-0.54 and PlaScope (reads) r_s_=-0.56) (**Supplementary Figure 8**). Again, plasmids in excess of 100kbp showed high recall variability while short plasmids (<20kbp) were generally fully recovered. There were some exceptions where plasmids <20kbp in length (n=578) had recall <0.05 for three tools: 90 plasmids (15.6%) with MOB-suite, 51 (8.9%) with PlaScope (assembly) and 50 (8.7%) with PlaScope (reads).

Comparing carriage of AMR genes across all plasmids in the isolates to F1 score showed no significant correlation (**Supplementary Figure 9**), with isolates carrying no AMR plasmids displaying a high degree of variability. Isolating individual plasmids and their individual AMR gene carriage confirmed that there was a minimal correlation between AMR genes and individual plasmid recall (MOB-suite r_s_=-0.15, PlaScope (assembly) r_s_=-0.27 and PlaScope (reads) r_s_=-0.27) (**Supplementary Figure 10**).

Contrastingly, the presence of IS elements appear to impact the performance of plasmid tools (**Supplementary Figure 11**). A correlation was most evident between the number of IS elements on plasmid recall, whereby a higher count of IS elements limited full plasmid recovery (MOB-suite r_s_=-0.5, PlaScope (assembly) r_s_=-0.67 and PlaScope (reads) r_s_=-0.68). Plasmids that carry >30 IS elements limited recall to less than 0.9191 for MOB-suite and 0.9263 for PlaScope (assembly). Using Unicycler assemblies as input resulted in a recall of ≤0.8815 for MOB-suite. However, of the 1,338 plasmids with less than 30 IS elements there were 522 (41.2%) plasmids not fully recovered (recall <0.95) by MOB-suite, 471 (35.1%) plasmids not recovered by PlaScope (assembly) and 448 (33.4%) plasmids not recovered by PlaScope (reads).

Lastly, we explored whether the presence of ‘shared’ sequences common to both plasmids and chromosome could impact performance (**Supplementary Figure 12**). Sharing of *k*-mers between plasmid and chromosome was calculated with Mash using a sketch size of 100,000.

We observed a similar, although less pronounced, trend to that of IS elements whereby a high level of *k*-mer sharing between plasmid and chromosome curtailed tool performance. This was most apparent at the level of individual plasmids (MOB-suite r_s_=-0.49, PlaScope (assembly) r_s_= −0.65 and PlaScope (reads) r_s_=-0.66) compared to total plasmid content (plasmidome) (MOB-suite r_s_=-0.26, PlaScope (assembly) r_s_=-0.29 and PlaScope (reads) r_s_=-0.31) (**Supplementary Figure 13**). Sharing of over 200 *k*-mers between the plasmidome and chromosome occurred in 89 isolates and limited the plasmidome F1 score for MOB-suite to 0.9851, for PlaScope (assembly) to 0.990641 and for PlaScope (reads) to 0.9943. Sharing of over 200 *k*-mers between individual plasmids and chromosome similarly limited individual plasmid recall scores to 0.9963 for MOB-suite, 0.9797 for PlaScope (assembly) and 0.9797 for PlaScope (reads). However, as we had observed for IS elements, there were exceptions where isolates with fewer than 200 shared *k*-mers still yielded poor plasmid recovery. There were 1,294 plasmids with less than 200 shared *k*-mers, of these 478 (36.9%) were not fully recovered (recall <0.95) by MOB-suite, 416 (32.1%) not recovered by PlaScope (assembly) and 395 (30.5%) not recovered by PlaScope (reads). This indicates complexities in resolving plasmid structures that cannot be explained by high IS counts and/or shared sequences.

For all metrics assessed, numerous plasmids exhibited poor performance despite not carrying IS elements or AMR genes and sharing no *k*-mers with the chromosome. There were 606 individual plasmids meeting these criteria, of which 37 had a recall <0.05 with both MOB-suite and PlaScope (both input modes). These plasmids may not be well represented in the underlying databases that the tools use for identification, as has been reported by previous studies (18). Databases used by these tools have largely skewed towards clinical isolates (18). To check this, we extracted the 37 plasmid sequences from the complete genomes and searched for matches in the MOB-suite and PlaScope databases using BLASTn with default settings. There were matches for 30 of the 37 plasmids in the MOB-suite database and for 28 plasmids in the PlaScope database. However, only two plasmids had matches covering more than 50% of their sequence across both databases, with the remaining plasmids only having low sequence coverage (<50%) (**Supplementary Table 5**). This indicates that poor performance of these plasmids is due to poor representation in tool databases and highlights the possibility that novel emergent plasmids will not be detected.

### Impact of data quality on performance

As the correlations between plasmid characteristics and tool performance we observed showed a high degree of variability, we further examined whether the quality and features of input data could affect performance. We therefore investigated the sequencing depth of plasmids, number of contigs and auNGA in input assemblies, and number of dead ends in input assembly graphs. Sequencing depth of the plasmids was calculated by mapping short reads to the hybrid assemblies and was normalised to the depth of chromosomal sequencing, whereby a depth of 1 represents the same sequencing depth shared between a plasmid and chromosome. Values above 1 indicate higher sequencing depth of a plasmid relative to the chromosome and is likely to occur in high copy number plasmids. Conversely, values less than 1 mean depth less than the chromosome, possibly occurring due to plasmid loss from some fraction of the sequenced population.

Examining individual plasmid recall scores against the normalised depth revealed that high sequencing depth resulted in complete plasmid recovery from read-based tools (PlaScope read input mode and plasmidSPAdes, **Supplementary Figure 14**) (PlaScope (reads) r_s_=0.46, plasmidSPAdes r_s_=0.5). Plasmid sequencing depth more than double that of the chromosome was sufficient to achieve complete recall for 72.8 % of plasmids for PlaScope (reads) and 74.0% for plasmidSPAdes. Even with normalized depth above 2 there were still 41 plasmids completely absent (recall of 0) for PlaScope (reads) and 60 for plasmidSPAdes. Sequencing depth of 1 or less resulted in more variable performance. When considering the impact of input genome assembly quality, we found only modest correlations between the number of contigs in the input assembly and tool F1 score, with fewer contigs associated with improved performance (MOB-suite r_s_=-0.21, PlaScope (assembly) r_s_=-0.29) (**Supplementary Figure 15**). We observed a stronger correlation between assembly quality (auNGA) and tool F1 score, with better quality (higher auNGA) equating to improved performance (MOB-suite r_s_=-0.41, PlaScope (assembly) r_s_=-0.41) (**Supplementary Figure 16**). We found no significant correlation between tool performance and number of dead ends in the assembly graph (**Supplementary Figure 17)**.

The expanding number of tools underscores the complexity of plasmid assembly while adding confusion for end-users when deciding which tool to use. Overall, no tool was able to recover the complete collection of plasmids, in part driven by short contigs in the genome assemblies likely causing loss of plasmid sequences. PlaScope achieved the highest overall performance, but key limitations for wider usage include its species-specific databases that include only *E. coli* and *K. pneumoniae.* Moreover, it is only capable of accepting genome assemblies produced by SPAdes, while a code update that allows Unicycler assembly input has not yet been packaged into conda. The second-best performing tool across all evaluation metrics, MOB-suite, offers broader species applicability which expands its utility amongst users. Further, its improved runtime and RAM usage metrics compared to PlaScope is another important consideration in analysis of large-scale datasets.

None of the tools benchmarked here attempt to recover complete plasmid sequences by following reference-matching paths through short-read assembly graphs. Some graph-based tools exist but address different tasks; Recycler finds cycles in the graph (*de novo*, not reference-guided), while PlasmidEC and gplas2 classify and bin contigs using graph links, yielding contig sets rather than a single path. These tools had been listed in our initial screening but were excluded from testing as they did not meet the inclusion criteria (e.g., Recycler could not be installed, PlasmidEC is specific to *E. coli* and, gplas2 requires an additional input file containing a binary plasmid-or-chromosomal prediction generated by an external classifier).

HyAsP also traces its own path through the assembly graph but uses a gene-based approach for this rather than full reference plasmid sequences; our results showed that this approach performed poorly. A full plasmid sequence reference-guided path-tracing approach could be effective, especially for large plasmids which are typically fragmented in assemblies: if a plasmid in the database (or a close relative) is present in the sample, its full sequence could be recovered by finding a matching path through the graph. This method has been applied manually using Bandage in previous studies to resolve plasmid sequences (51) but no existing software performs this automatically. An automated implementation could potentially outperform current tools, particularly in outbreak settings (e.g. tracking AMR or virulence plasmids in health-care settings) where a close-match reference plasmid from hybrid assembly is available.

## CONCLUSION

The critical role and clinical relevance of plasmids, particularly in the dissemination of AMR and virulence in nosocomial pathogens like *K. pneumoniae*, underpins the need for robust and scalable tools for plasmid identification and reconstruction from large-scale short-read datasets. As yet, no tool offers a comprehensive and universally reliable solution to assembling plasmids from short-read sequence data. Our benchmarking identified PlaScope and MOB-suite as the best tools, although the former can only accept unmodified genome assemblies from SPAdes. Significant advances in this space will open-up new avenues in both public health surveillance and exploring the evolution and movement of plasmids. Novel tools developed in this space should offer substantial advancements over existing tools and be comprehensively evaluated using large, standardised bacterial datasets with diverse plasmid content to ensure consistency across benchmarking studies.

## Supporting information

All tables

## Data Availability

The Nextflow pipeline and outputs with code for producing figures is available at: github.com/C-Connor/PlasmidToolBenchmarking. The *Klebsiella* data from Hetland *et al.* (30) are available at BioProject: PRJEB74192. Associated metadata of plasmids are available as supplementary files from Winkler *et al.* (31)

## Author contributions

Conceptualization: CHC, CLG, DJI, MMCL Data Curation: CHC, MAW, MMCL

Formal analysis: CHC, RRW Funding Acquisition: DJI, MMCL

Methodology: CHC, RRW, CLG, DJI, MMCL Resources: MAW, IHR

Supervision: DJI

Writing - original draft preparation: CHC

Writing - reviewing & editing: CHC, RRW, CLG, MAW, IHR, DJI, MMCL All authors reviewed and approved the final manuscript.

